# Tyrosine hydroxylase deletion in *Ntsr1* neurons confers resistance to diet-induced weight gain: an energy-balance analysis

**DOI:** 10.64898/2026.11.05.764678

**Authors:** Renqi Wu, Alyssa Gomez, Justine T. Wong, Ruth Gordillo, Anna K. Kamitakahara, Andrew D. Steele

## Abstract

High-fat diet (HFD) disrupts circadian feeding rhythms and promotes diet-induced obesity (DIO), but the neural mechanisms linking diet to energy balance remain unclear. Previously, we demonstrated that dopamine production in neurotensin receptor 1 (*Ntsr1*)-expressing neurons was required for HFD-induced daytime overfeeding and DIO. Here, we tested whether DIO resistance in *Ntsr1^Cre^;Th^flox^* conditional knockout (cKO) mice was explained by altered energy expenditure. We replicated resistance to HFD-induced weight gain and assessed food intake, indirect calorimetry, activity, body temperature, fecal energy, fecal nonesterified fatty acids, and the gut microbiome. Indirect calorimetry on standard chow diet (SCD) was confounded by reduced intake and body weight loss, whereas HFD-fed cKO mice consumed food readily and resisted weight gain. Unlike our previous study, daytime feeding was not suppressed; instead, cKO mice were slightly and variably hypophagic on HFD but not SCD. DIO resistance was not accompanied by increased energy expenditure, activity, or body temperature. Rather, cKO mice showed reduced energy expenditure and activity and lower body temperature on both diets. In the replication cohort, HFD fecal output was lower in cKO mice, whereas the independent bomb-calorimetry cohort showed no genotype difference in fecal mass; fecal energy density and nonesterified fatty acid concentration were unchanged. Preliminary analysis of the microbiome was consistent with an obesity-resistant gut profile. Together, these findings indicate that the feeding phenotype is diet- and experiment-dependent and that DIO resistance is not explained by increased energy expenditure, activity, thermogenesis, or detectable changes in fecal energy or fat content.

## Introduction

First isolated from bovine hypothalamus in 1973, the 13-amino-acid peptide neurotensin (Nts) has occupied a distinctive place in neuroscience as a neuropeptide linking internal physiological state with motivated behavior [1, 2]. Its widespread distribution, actions on dopamine systems, and signaling through neurotensin receptor 1 (Ntsr1) and neurotensin receptor 2 (Ntsr2) have implicated it in reward, addiction, thermoregulation, nociception, locomotion, and feeding [2–6]. Of particular relevance to the present study, work from Leinninger and colleagues and others has established neurotensin as an important regulator of feeding and energy balance: neurotensin produced in the brain and gut engages hypothalamic, brainstem, gastrointestinal, and mesolimbic pathways to influence meal termination, food reward, physical activity, energy expenditure, and body weight [4, 7]. Studies of lateral hypothalamic neurotensin neurons and VTA Ntsr1 neurons have shown how neurotensin pathways can connect leptin-related metabolic signals to dopamine-dependent motivation and body-weight regulation [6, 8]. Additional work highlights that neurotensin receptor signaling in mesolimbic circuits directly modulates synaptic transmission and feeding responses, reinforcing the functional interdependence of neurotensin and dopamine systems in energy homeostasis [3, 9]. Importantly, neurotensin is neither uniformly anorexigenic nor uniformly obesogenic; its effects vary with the receptor, cell population, projection target, diet, and physiological state [4, 5, 10]. Understanding the circuit- and state-dependence of neurotensin signaling using conditional genetic approaches for anatomical restriction will be critical to understanding how metabolic need and reward interact [11].

Disruption of circadian feeding rhythms is increasingly recognized as a contributor to obesity and metabolic disease [12, 13]. In nocturnal rodents, standard chow diet (SCD) intake is normally concentrated during the dark, active phase, whereas high-fat diet (HFD) exposure increases food consumption during the light, rest phase [14, 15]. This redistribution of calories across the circadian cycle promotes weight gain by blunting rhythmic metabolic gene expression and altering adipocyte thermogenic mechanisms [16, 17]. Thus, understanding how HFD engages neural circuits that override normal circadian control of feeding may reveal mechanisms of obesity that are not apparent from total caloric intake alone.

Dopamine neurons in the ventral midbrain are well positioned to link HFD to altered circadian feeding behavior [18]. Although VTA dopamine neurons are classically studied for their projections to mesolimbic reward centers, they also provide functional input to D1R-expressing neurons in the suprachiasmatic nucleus (SCN), the central circadian pacemaker, where they mediate shifting of activity cycles in response to simulated jet lag [19–22]. Interestingly, global deletion of dopamine D1 receptor leads to a profound resistance to diet-induced obesity (DIO) [15]. Remarkably, restoration of D1R to the nucleus accumbens (NAc) did not confer susceptibility to DIO but restoration to the SCN did [15]. These findings expanded our understanding of the SCN beyond its role as a circadian pacemaker, identifying it as an integrative node that links dopamine signaling and circadian timing to feeding behavior and susceptibility to DIO. More broadly, they suggest that dopamine input to the SCN may provide a mechanism by which the motivational value of HFD is coupled to the temporal organization of feeding and long-term energy balance.

In follow-up studies, we determined that deletion of *tyrosine hydroxylase* (*Th*) from *Ntsr1*-expressing neurons prevents DIO, possibly by blunting daytime feeding, while viral restoration of TH in the VTA restores both DIO and day-eating of HFD [23]. These findings support a model in which *Ntsr1*-expressing dopamine neurons promote obesity by disrupting circadian regulation of meal timing. However, the physiological basis of this DIO resistance remains unresolved. Loss of dopamine production in *Ntsr1* neurons could protect against weight gain by increasing thermogenesis, elevating whole-animal energy expenditure, reducing nutrient (especially fat) uptake, or by altering microbiome composition. Distinguishing among these possibilities is important because each implies a different mechanism by which a genetically defined dopamine circuit contributes to obesity.

Here, we investigated the mechanism(s) underlying resistance to DIO in *Ntsr1^Cre^*;*Th^flox^* cKO male mice. We combined replication studies of body weight and day/night food intake with additional measures that included indirect calorimetry, activity measurements, and temperature monitoring to determine whether the phenotype is explained by altered energy expenditure or thermogenesis. We also measured fecal energy content and lipid composition to test whether altered nutrient absorption [24, 25] may contribute to the DIO resistance of *Ntsr1^Cre^*;*Th^flox^*cKO male mice. Lastly, we examined microbiome composition to test whether this may alter the susceptibility to DIO [26, 27].

## Methods

### Mouse strains, husbandry, and cohorts

The experiments described herein were approved by the California State Polytechnic, Pomona Institutional Animal Care and Use Committee (IACUC) under protocols: 23.006, 23.009, 25.018, and 26.045.

Mice were single housed in static microisolator cages containing Sani-Chip bedding (Envigo 7090), a plastic house, and a cotton nestlet under a 12 h light/dark cycle with *ad libitum* access to food and water. Housing conditions were maintained at an ambient temperature of 21-24 °C and relative humidity between 20–45%. Male mice were used for all experiments reported here. Adult mice (≥10 weeks of age) were used for all analyses.

Genomic DNA was obtained from tail biopsies collected from mice at 12–14 days of age. Samples were submitted to Transnetyx (Cordova, TN) for analysis. Genotyping was performed using real-time PCR to detect the presence of *Cre* recombinase, *Th^flox^* allele, and *Th^Wt^* alleles.

The *Ntsr1^Cre^* line was obtained from the breeding colony of Prof. Gina Leinninger (Michigan State University) and is available via the Jackson Laboratory (Bar Harbor, ME) under stock number 033365. The *Th^flox^* allele, containing loxP sites flanking the first exon of *Th*, was obtained from Dr. Richard Palmiter (University of Washington)[28]. Experimental *Ntsr1^Cre^*;*Th^flox^*cKO mice were Cre-positive, carried the *Th^flox^* allele, and lacked a wild-type *Th* allele (*Th*^flox/flox^). Littermate control mice were *Ntsr1^Cre^*-negative or *Ntsr1^Cre^*-positive and carried at least one *Th^Wt^* allele (*Th^flox/Wt^*, *Th*^Wt/Wt^).

Different assays were performed in different animal cohorts, as follows (Supplementary Figure 1). The replication cohort (*n* = 10 control and *n* = 7 cKO males) was used for body weight and food intake (Figure 1A–F) and for home-cage fecal output (Figure 4A–D). Fecal NEFA concentrations (Figure 4I, J; n = 8 control and *n* = 6 cKO) and microbiome composition (Figure 5; *n* = 7 control and *n* = 3 cKO) were measured in subsets of this replication cohort. For Figure 1G–N, *n* = 6 control and *n* = 4 cKO. For Figures 2 and 3, *n* = 10 control and *n* = 6 cKO males were used for metabolic measurements. For Figure 4E, *n* = 13 control and *n* = 10 cKO samples were analyzed, and for Figure 4F, *n* = 14 control and *n* = 9 cKO. Home-cage food intake in the bomb calorimetry cohort (Figure 4G, H) was measured in *n* = 7 control and *n* = 7 cKO males.

**Figure 1.**
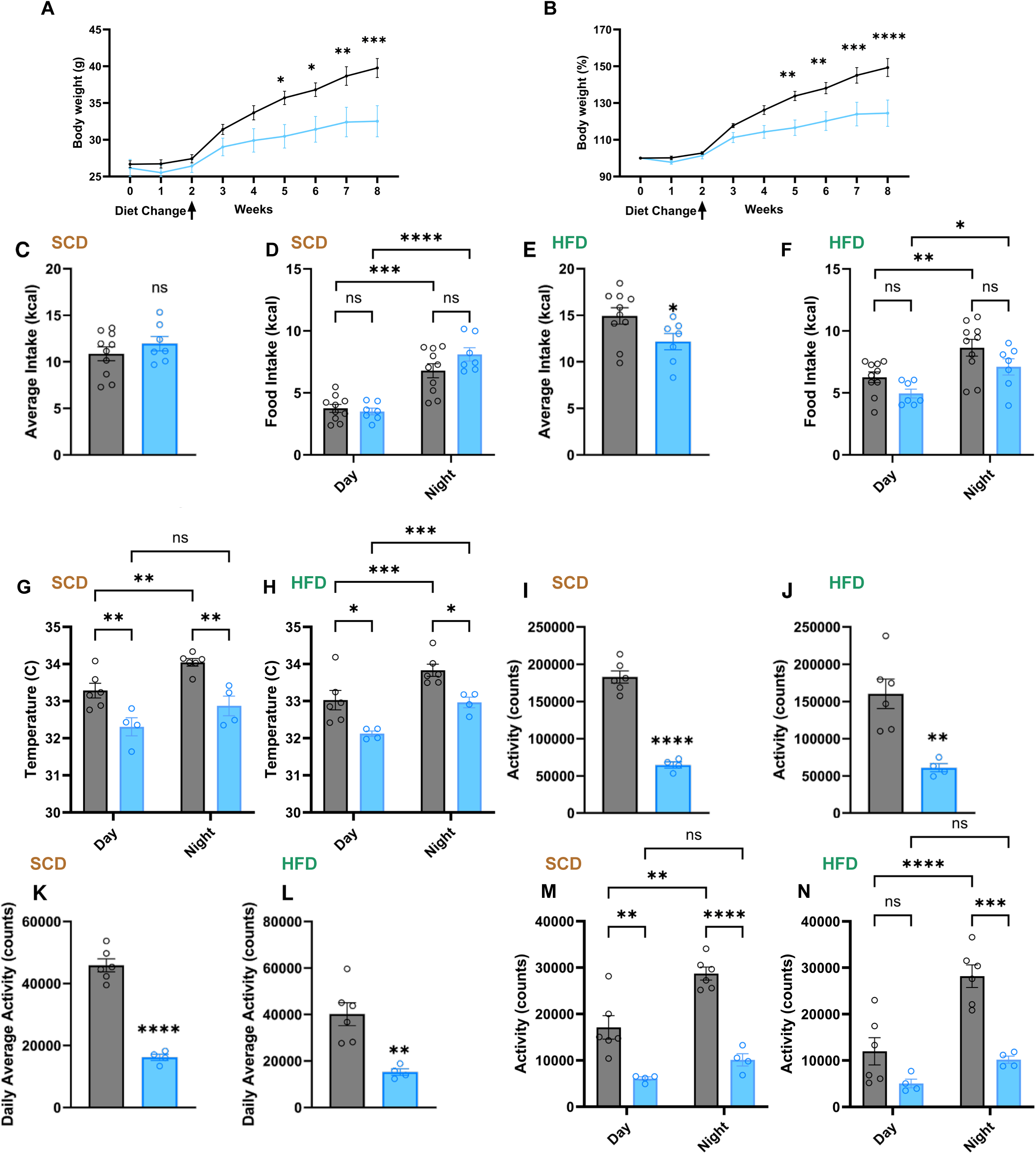
Body-weight, food intake, locomotor activity, and body temperature in. (A, B) Body-weight in grams and as percentage of baseline (week 0), respectively, measured weekly from week 0 through week 8; the diet change from SCD to HFD occurred after week 2. Control *n* = 10 and cKO *n* = 7. (C–F) Average daily food intake and day/night food intake on SCD (C, D) and at week 5 of HFD (E, F); control *n* = 10, cKO *n* = 7. (G, H) Day/night subcutaneous body temperature measured with implanted Nanotags under SCD (G) and HFD (H). (I–N) Nanotag total activity (I, J), daily average activity (K, L), and day/night activity (M, N) under SCD and HFD. Nanotag measures: control *n* = 6, cKO *n* = 4. Bars and points show mean ± SEM and individual mice, respectively. Gray denotes control and blue denotes cKO. ns, *p* ≥ 0.05; \**p* < 0.05; \*\**p* < 0.01; \*\*\**p* < 0.001; \*\*\*\**p* < 0.0001.

**Figure 2.**
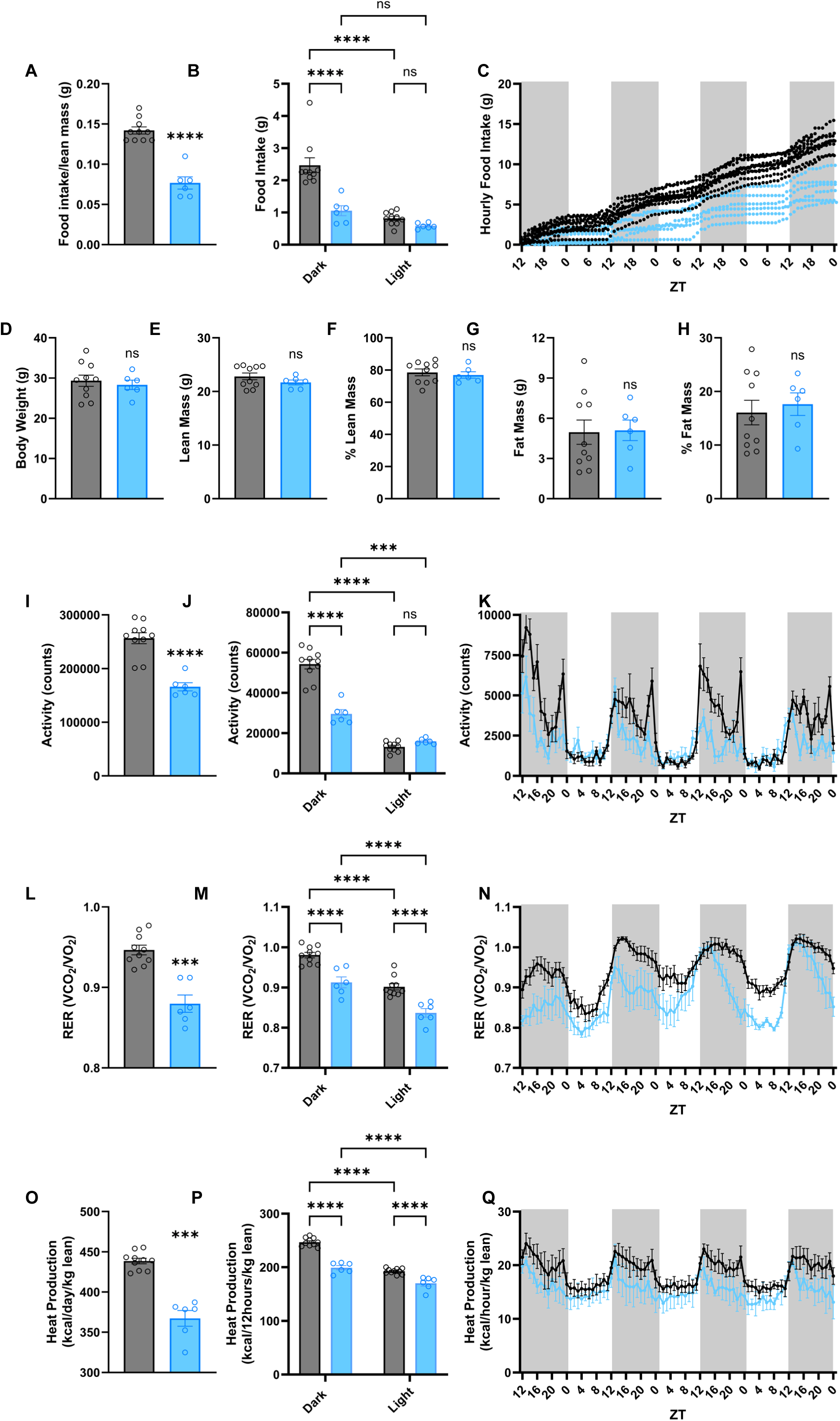
Body composition, locomotor activity, respiratory exchange ratio, heat production, and food intake during standard-chow indirect calorimetry. Mice maintained on SCD were assessed in metabolic cages. (A-C) Daily food intake normalized to lean mass, average dark- and light-phase food intake, and cumulative food intake. (D-H) Body weight, lean mass, percentage lean mass, fat mass, and percentage fat mass, respectively. (I-K) Total locomotor activity, average dark- and light-phase activity, and hourly activity. (L-N) Mean respiratory exchange ratio (RER; VCO_₂_/VO_₂_), average dark- and light-phase RER, and hourly RER. (O-Q) Daily mean heat production normalized to lean mass, average dark- and light-phase heat production, and hourly heat production. Control *n* = 10; cKO *n* = 6. Hourly and cumulative plots are shown descriptively. Gray indicates control and blue indicates cKO; gray background shading denotes the dark phase. Bars and points show mean ± SEM and individual mice, respectively; time courses show mean ± SEM. ns, *p* ≥ 0.05; \**p* < 0.05; \*\**p* < 0.01; \*\*\**p* < 0.001; \*\*\*\**p* < 0.0001.

**Figure 3.**
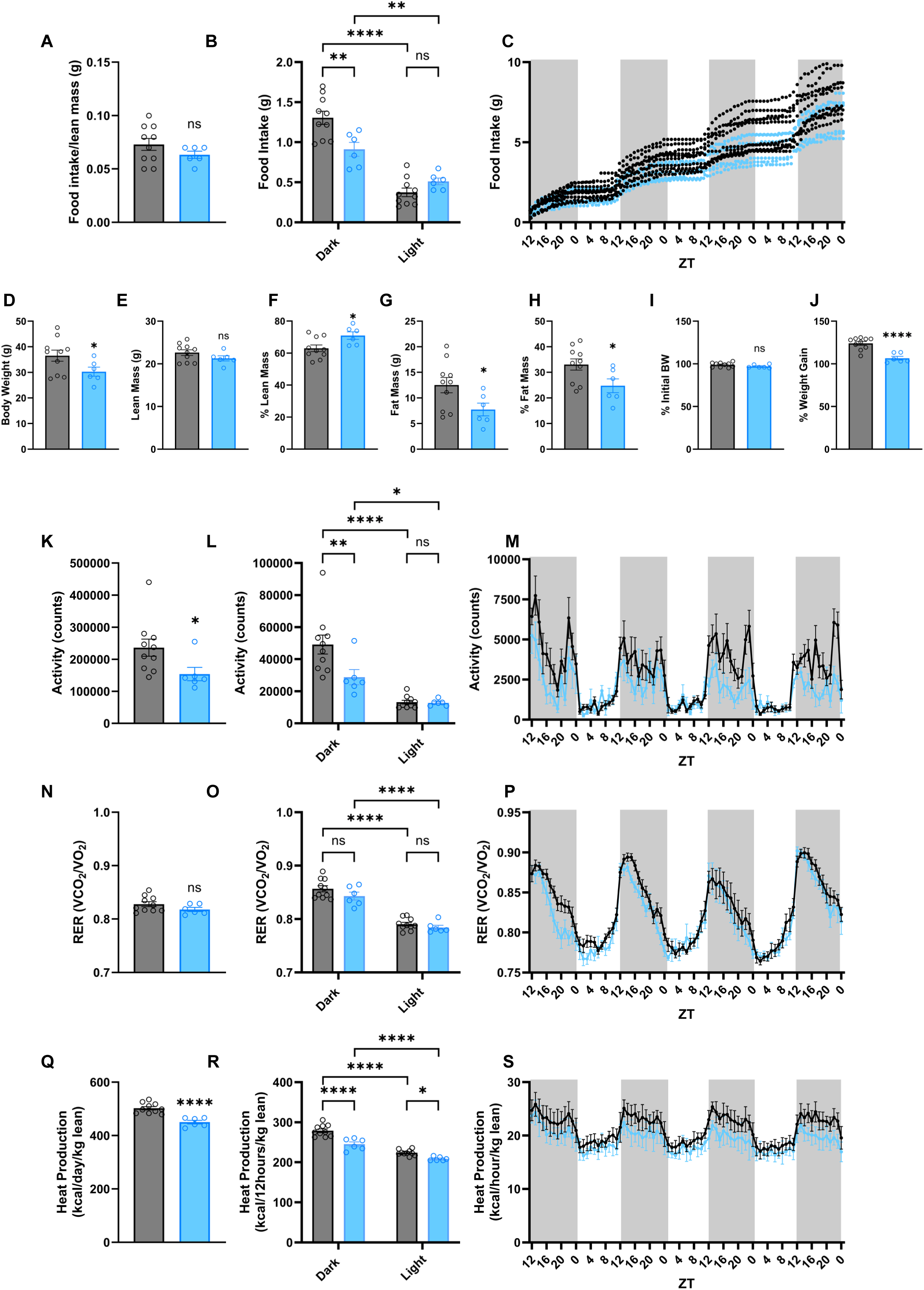
Body composition, locomotor activity, respiratory exchange ratio, heat production, and food intake during high-fat-diet indirect calorimetry. The same mice shown in Figure 2 were assessed after 3 weeks of HFD exposure (study week 5). (A-C) Daily food intake normalized to lean mass, average dark- and light-phase food intake, and cumulative food intake. (D-J) Body weight, lean mass, percentage lean mass, fat mass, percentage fat mass, body weight after versus before the metabolic-cage recording period expressed relative to initial body weight, and body-weight gain relative to the start of HFD exposure, respectively. (K-M) Total locomotor activity, average dark- and light-phase activity, and hourly activity. (N-P) Mean RER, average dark- and light-phase RER, and hourly RER. (Q-S) Daily mean heat production normalized to lean mass, average dark- and light-phase heat production, and hourly heat production. Control *n* = 10; cKO *n* = 6. Hourly and cumulative plots are shown descriptively. Gray indicates control and blue indicates cKO; gray background shading denotes the dark phase. Bars and points show mean ± SEM and individual mice, respectively; time courses show mean ± SEM. ns, *p* ≥ 0.05; \**p* < 0.05; \*\**p* < 0.01; \*\*\**p* < 0.001; \*\*\*\**p* < 0.0001.

**Figure 4.**
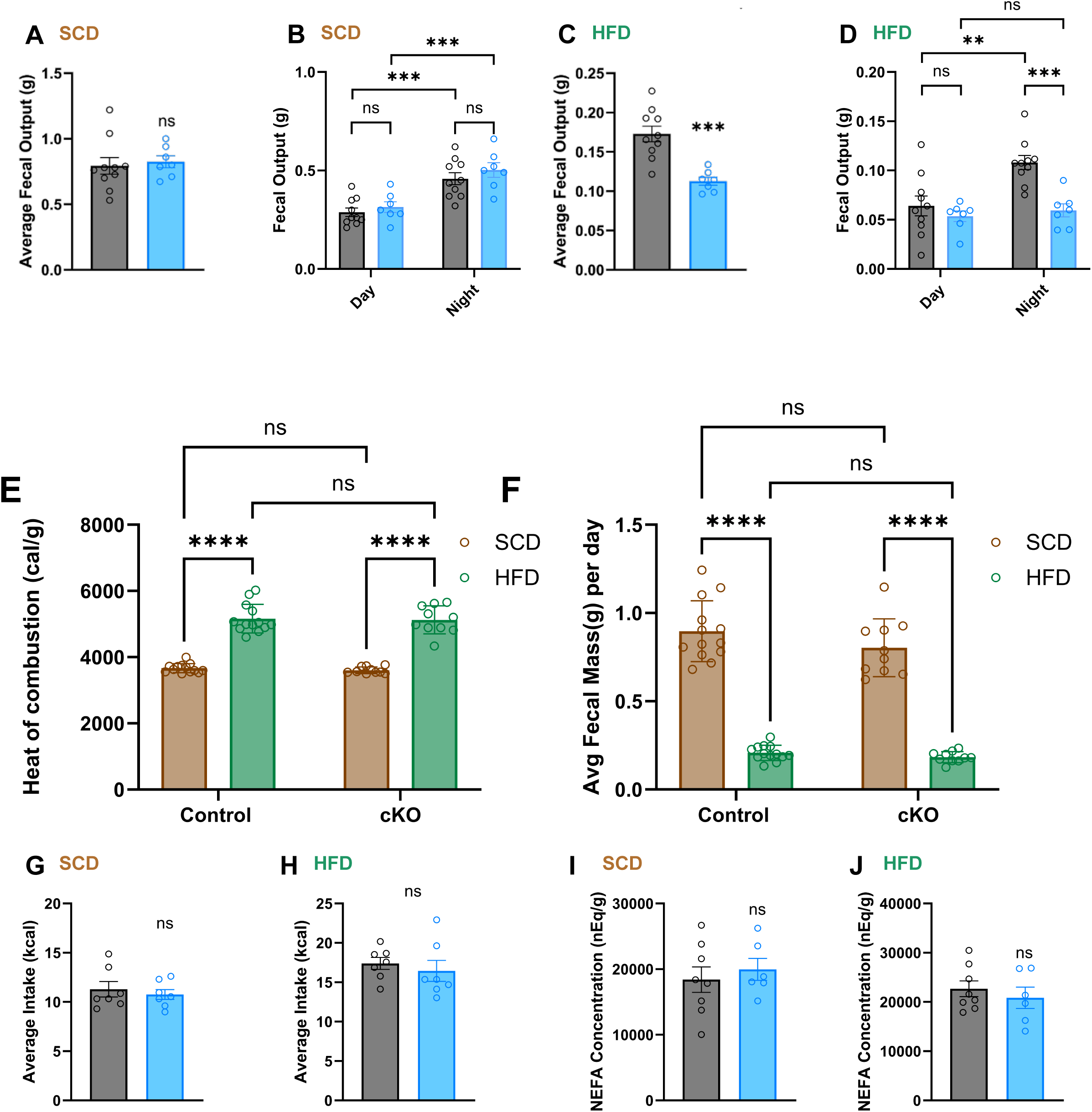
Fecal output, fecal energy density, food intake, and nonesterified fatty acids. (A–D) Fecal output measured in the home cage: average daily fecal output and day/night fecal output on SCD (A, B) and HFD (C, D); control *n* = 10, cKO *n* = 7. (E, F) Heat of combustion of dried fecal material measured by bomb calorimetry, and average fecal mass produced per day under SCD and HFD conditions. Feces were collected every 24 h; heat of combustion, control *n* = 13 and cKO *n* = 10 samples per diet (E); fecal mass per day, control *n* = 14 and cKO *n* = 9 (F). (G, H) Average daily home-cage food intake on SCD (G) and HFD (H) in the bomb calorimetry cohort; control *n* = 7, cKO *n* = 7. (I, J) Fecal nonesterified fatty acid (NEFA) concentration on SCD (I) and HFD (J); control *n* = 8, cKO *n* = 6. Heat of combustion was analyzed by two-way ANOVA (genotype × diet) with Tukey-adjusted comparisons; single-value summaries were analyzed using two-sided unpaired Welch’s *t* tests and dark/light data by two-way repeated-measures ANOVA (genotype × phase) with Šídák-adjusted comparisons between genotypes within each phase and between phases within each genotype. Gray denotes control and blue denotes cKO; bars and points show mean ± SEM and individual mice, respectively. ns, *p* ≥ 0.05; \**p* < 0.05; \*\**p* < 0.01; \*\*\**p* < 0.001; \*\*\*\**p* < 0.0001.

**Figure 5.**
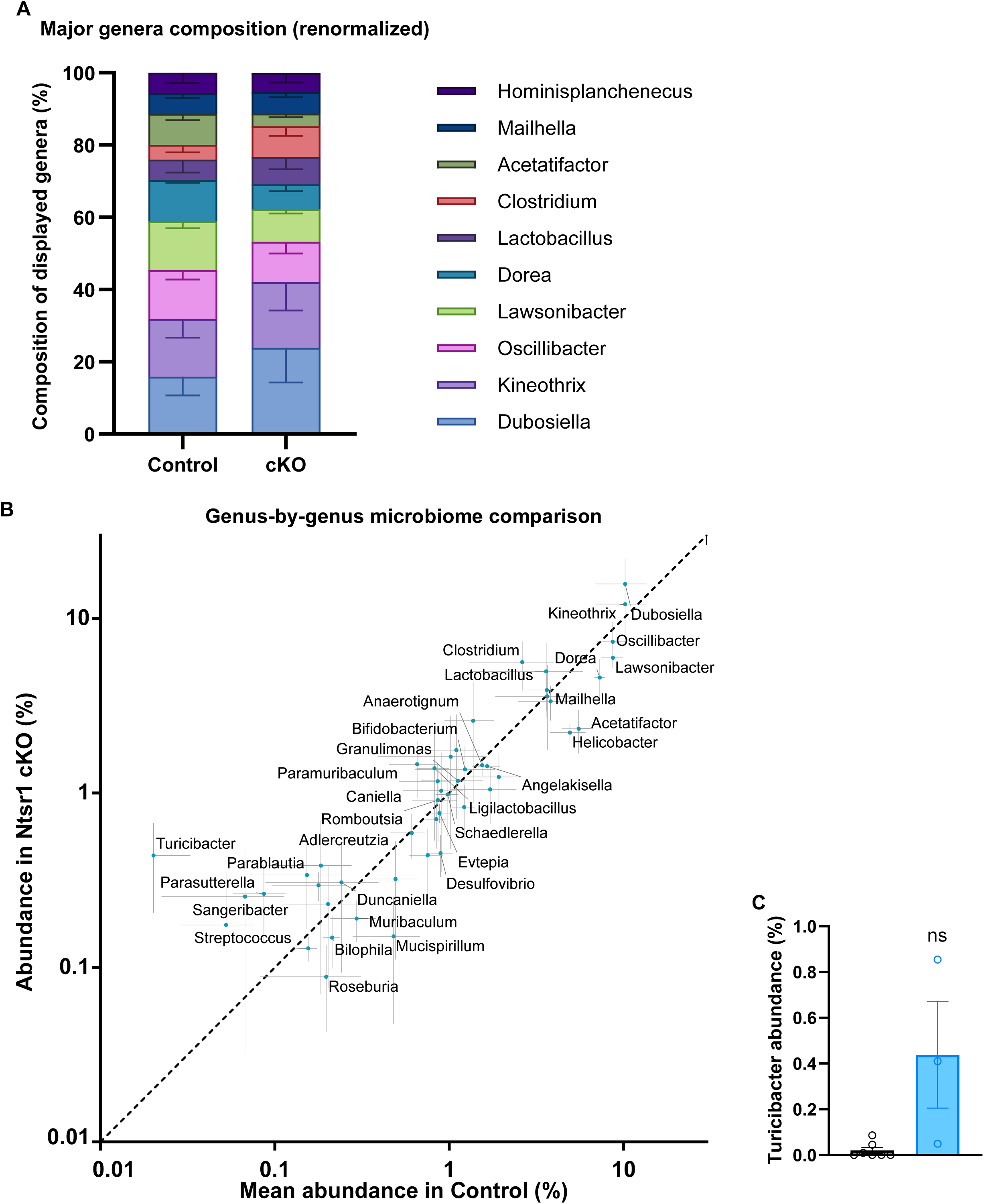
Exploratory fecal microbiome composition in *Ntsr1-TH-cKO* mice. (A) Renormalized relative abundance of the major genera in control (*n* = 7) and cKO (*n* = 3) mice. (B) Genus-by-genus comparison of mean abundance in cKO versus control mice, with the line of identity. Fecal microbiome profiling was performed by Transnetyx. Alpha diversity (Shannon index) and individual genera did not differ significantly between genotypes. (C) *Turicibacter* relative abundance in control (*n* = 7) and cKO (*n* = 3) mice (Welch’s t test, ns). Error bars show SEM.

### Diet and body-weight monitoring

Mice were maintained on standard chow diet (SCD; Teklad 2018, Envigo; 18% fat, 24% protein, and 58% carbohydrate by calories; 3.1 kcal/g) and weighed at baseline (week 0) and weekly thereafter. After the week 2 measurement, mice were switched to a high-fat diet (HFD; Bio-Serv F3282; 59% fat, 15% protein, and 26% carbohydrate by calories; 5.49 kcal/g) for 6 additional weeks. Body weight was expressed in grams and as a percentage of each mouse’s week 0 body weight.

### Home-cage food intake and fecal output

Food intake and fecal output were measured in home cages on SCD (study week 2) and after 3 weeks of HFD (study week 5). Food was provided in overhead wire racks, and food was weighed and feces were collected at lights-on (ZT0) and lights-off (ZT12), so that light- and dark-phase values could be obtained separately; spilled food on the cage floor was collected and included in the measurements. Food intake in grams was converted to kcal by multiplying by the energy density of the diet (SCD, 3.1 kcal/g; HFD, 5.49 kcal/g). In the bomb calorimetry cohort (Figure 4G, H), 40 g of food was provided and the remaining food was weighed after 3 days on SCD and 4 days on HFD; average daily intake was calculated as the amount eaten divided by the number of days and converted to kcal in the same way.

Feces were collected every 12 h to obtain separate light- and dark-phase output and stored at –80°C. Fecal nonesterified fatty acids (NEFA) were measured at the University of Texas Southwestern Medical Center (UTSW). For bomb calorimetry, feces were collected every 24 h and dried, and heat of combustion was measured.

For analysis of microbiome, fecal samples were collected immediately upon excretion, placed directly into the stabilization medium provided by Transnetyx (Cordova, TN), and submitted to Transnetyx, which performed sequencing and genus-level taxonomic profiling. Genus-level relative abundances were renormalized to the major genera shown.

### Subcutaneous temperature and activity using Nanotags

Body temperature and locomotor activity were recorded using subcutaneously implanted Nanotag devices (Stoelting), which were decontaminated with ethylene oxide gas before implantation; measurements were recorded at 5-minute intervals as done previously [29]. Mice were anesthetized with 3–5% isoflurane in O_₂_, maintained at 1–3% isoflurane with 1 L/min O_₂_, and stabilized in a stereotaxic frame (Kopf, Model 1900).

After ophthalmic ointment was applied and the surgical site was clipped and cleaned with chlorhexidine and 70% isopropyl alcohol, a 2.0–2.5 cm incision was made on the back, and a subcutaneous pocket was created by blunt dissection. The sterile Nanotag was inserted, the incision was closed with 9 mm EZ Clip wound clips (Braintree Scientific), and Marcaine (Hospira, 0.5%, 5 mg/mL; 0.1 mL) was applied to the incision site. Mice received ketoprofen (0.1 mL of 0.5 mg/mL, subcutaneously) at surgery and daily for the following 3 days, were weighed before and after surgery, and had wound clips removed 2 weeks after surgery. For each mouse, temperature and activity were averaged separately over the light and dark phases, and activity was also summed over the full recording (total activity) and averaged per day (daily average activity).

### Indirect calorimetry and body composition

Indirect calorimetry and body composition were measured at the Metabolic Phenotyping Core of The Saban Research Institute of Children’s Hospital Los Angeles. Body composition (fat and lean mass) was measured by quantitative magnetic resonance using an EchoMRI-700 before and after each metabolic-cage recording. Mice were individually housed in metabolic cages (TSE LabMaster/PhenoMaster, TSE Systems) for 3.5-day recordings on SCD (study week 1) and after 3 weeks of HFD (study week 5). Initial pilot testing revealed that cKO mice failed to drink from the metabolic cage lixits. To mitigate the effects of dehydration in cKO mice and provide equal housing conditions among all groups, a non-caloric hydration gel cup (HydroGel) was provided in all cages. Total body water content measured by EchoMRI at the start and end of each 3.5 day metabolic cage test period confirmed adequate hydration of all mice during testing. O_₂_ consumption, CO_₂_ production, food intake, and locomotor activity were recorded continuously. RER was calculated as VCO_₂_/VO_₂_, and heat production was calculated using the Weir equation and normalized to lean mass [30, 31]. Data were summarized hourly, for each light and dark period, and as daily averages or study totals. Body weight after each recording was expressed as a percentage of body weight before it.

### Statistical analysis

Statistical analyses were performed using GraphPad Prism 10.6. Data are presented as mean ± SEM, and *n* refers to individual mice. Two-group comparisons used two-sided unpaired *t* tests with Welch’s correction. Light/dark (or day/night) data, which were measured repeatedly in the same mice, were analyzed by two-way repeated-measures ANOVA with genotype as the between-subject factor and phase (dark versus light, averaged within each mouse) as the within-subject factor, followed by Šídák-adjusted comparisons between genotypes within each phase and between phases within each genotype (each set of two comparisons treated as one family). Body-weight time courses were analyzed by two-way repeated-measures ANOVA (genotype x week), followed by Šídák-adjusted comparisons between genotypes at each week. Hourly and cumulative time courses are presented descriptively. Microbiome alpha diversity (Shannon index) and the abundance of individual genera were compared between genotypes using unpaired t tests with Welch’s correction; p values across the 48 genera were corrected using the Benjamini-Hochberg false-discovery-rate method; relative abundances were renormalized after excluding unclassified reads. Statistical significance was defined as *p* < 0.05.

## Results

### Body weight, food intake, locomotor activity, and body temperature

We first monitored body weight in control and *Ntsr1^Cre^*;*Th^flox^*cKO male mice. Mice from three cohorts were entered into the study at 9–12 weeks of age and weighed at baseline (week 0) and weekly for two weeks while maintained on standard chow diet (SCD), yielding three measurements in total, after which they were switched to a high-fat diet (HFD) for 6 additional weeks. Between week 2 (the last SCD measurement) and week 5, control mice (*n* = 10) gained 8.3 ± 0.6 g and *Ntsr1^Cre^*;*Th^flox^*cKO mice (*n* = 7) gained 4.0 ± 1.0 g (mean ± SEM). Between week 2 and week 8, control mice (*n* = 10) gained 12.4 ± 1.2 g (approximately 2.1 g per week), whereas *Ntsr1^Cre^*;*Th^flox^* cKO mice (*n* = 7) gained 6.1 ± 1.6 g (approximately 1.0 g per week) (Figure 1A). For body weight in grams, a two-way repeated-measures ANOVA identified significant effects of genotype (*F(1,15) = 6.43, p = 0.0228),* week (*F*(8,120) = 81.14, p < 0.0001), and genotype x week (*F*(8,120) = 8.17, p < 0.0001; Figure 1A). Šídák-adjusted comparisons showed that cKO mice weighed significantly less than controls from week 5 onward (week 5 *p* = 0.0182, week 6 *p* = 0.0136, week 7 *p* = 0.0022, and week 8 p = 0.0002), with no significant genotype difference at weeks 0–4 (p ≥ 0.204). Normalized body weight, calculated relative to week 0, was analyzed using the same two-way repeated-measures ANOVA, which identified significant effects of genotype (*F*(1,15) = 10.70, *p* = 0.0052), week (*F*(8,120) = 74.96, *p* < 0.0001), and genotype x week (*F*(8,120) = 7.29, *p* < 0.0001; Figure 1B). Šídák-adjusted comparisons showed that cKO mice had gained significantly less weight than controls from week 5 onward (week 5 *p* = 0.0036, week 6 *p* = 0.0024, week 7 *p* = 0.0002, and week 8 *p* < 0.0001), with no significant genotype difference at weeks 0–4 (*p* ≥ 0.119).

Food intake on SCD and HFD was measured over three-day periods, with intake quantified separately during the light and dark phases. Average daily food intake on SCD was similar between control (*n*=10) and *Ntsr1^Cre^*;*Th^flox^*cKO (*n*=7) mice (Welch’s *t*(14.10) = 1.01, *p* = 0.328; Figure 1C). Two-way repeated-measures ANOVA (genotype × phase) of day versus night intake identified a significant phase effect (*F*(1,15) = 74.16, *p* < 0.0001), with no genotype effect (*F*(1,15) = 1.21, *p* = 0.289) or genotype × phase interaction (*F*(1,15) = 3.12, *p* = 0.0977; Figure 1D), and control and cKO mice did not differ within the day or night phase (Šídák-adjusted *p* = 0.912 and *p* = 0.106, respectively). At week 5, average daily HFD intake was lower in cKO mice (Welch’s *t*(14.38) = 2.22, *p* = 0.0431; Figure 1E). The day/night analysis identified effects of *p*hase (*F*(1,15) = 22.10, *p* = 0.0003) and genotype (*F*(1,15) = 4.72, *p* = 0.0462), without a genotype × phase interaction (*F*(1,15) = 0.060, *p* = 0.810; Figure 1F), and the genotypes did not differ significantly within either phase (day *p* = 0.223; night *p* = 0.131).

To assess the effects of diet on body temperature and activity, we implanted subcutaneous Nanotags in control mice (*n* = 6) and *Ntsr1^Cre^*;*Th^flox^*cKO (*n* = 4) and followed the same animals longitudinally as they transitioned from normal chow to HFD. Temperature and activity were recorded continuously for approximately 4 days on each diet, and the day and night values reported are averages across that recording. Under SCD, two-way repeated-measures ANOVA (genotype × phase) identified effects of phase (*F*(1,8) = 21.60, *p* = 0.00165) and genotype (*F*(1,8) = 19.70, *p* = 0.00217), with no genotype × phase interaction (*F*(1,8) = 0.477, *p* = 0.509; Figure 1G). cKO mice had lower temperatures during both the day (Šídák-adjusted *p* = 0.00623) and night (*p* = 0.00143). Under HFD, temperature again showed effects of *p*hase (*F*(1,8) = 81.65, *p* < 0.0001) and genotype (*F*(1,8) = 10.41, *p* = 0.0121), with no interaction (*F*(1,8) = 0.035, *p*= 0.857; Figure 1H). cKO temperature was lower during both day and night (*p* = 0.0131 and *p* = 0.0167, respectively, Figure 1H).

Nanotag locomotor activity was also reduced in cKO mice. Total activity over the SCD recording period was lower in cKO mice (Welch’s *t*(7.10) = 12.82, *p* < 0.0001; Figure 1I), as was total activity during HFD (Welch’s *t*(5.75) = 4.87, *p* = 0.00315; Figure 1J). Daily average activity was similarly lower on SCD (Welch’s *t*(7.06) = 12.73, *p* < 0.0001; Figure 1K) and HFD (Welch’s *t*(5.78) = 4.86, *p* = 0.00312; Figure 1L). SCD day/night activity showed effects of phase (*F*(1,8) = 11.71, *p* = 0.00905) and genotype (*F*(1,8) = 116.76, *p* < 0.0001), while the genotype × phase interaction was not significant (*F*(1,8) = 2.70, *p* = 0.139; Figure 1M); cKO activity was lower during both the day (Šídák-adjusted *p* = 0.00152) and night (*p* < 0.0001). During HFD, day/night activity showed effects of phase (*F*(1,8) = 53.29, *p* < 0.0001), genotype (*F*(1,8) = 15.92, *p* = 0.00400), and genotype × phase interaction (*F*(1,8) = 14.40, *p* = 0.00528; Figure 1N), reflecting lower cKO activity at night (*p* = 0.0002) but not during the day (*p* = 0.120).

### Indirect calorimetry during standard chow diet

Daily food intake normalized to lean mass was lower in cKO mice (Welch’s *t*(8.42) = 7.43, *p* < 0.0001; Figure 2A). Dark/light food intake showed effects of phase (*F*(1,14) = 36.33, *p* < 0.0001), genotype (*F*(1,14) = 25.37, *p* = 0.0002), and genotype × *p*hase (*F*(1,14) = 11.18, *p* = 0.0048; Figure 2B); cKO mice ate less during the dark phase (Šídák-adjusted *p* < 0.0001) but not the light phase (*p* = 0.559). Cumulative food intake across the 3.5-day metabolic-cage recording is shown in Figure 2C.

Mice did not eat normally after transferring to the metabolic cages on SCD. Over the 3.5-day recording, cKO mice ate less chow per gram of lean mass than controls and lost more body weight: post-recording body weight was 89.2 ± 0.9% of pre-recording weight in cKO mice versus 98.8 ± 0.6% in controls (Figure 2A, Welch’s *t*(9.92) = 8.63, *p*< 0.0001). The SCD calorimetry data should therefore be interpreted in light of this reduced intake. We next assessed body composition and metabolic measures in control and cKO mice during a 3.5-day SCD metabolic-cage recording. At the start of the recording, body weight did not differ between genotypes (Welch’s *t*(13.87) = 0.56, *p* = 0.583; Figure 2D). Lean mass (Welch’s *t*(13.98) = 1.46, *p* = 0.165; Figure 2E), percentage lean mass (Welch’s *t*(13.27) = 0.56, *p* = 0.583; Figure 2F), fat mass (Welch’s *t*(13.80) = 0.12, *p* = 0.906; Figure 2G), and percentage fat mass (Welch’s *t*(13.41) = 0.50, *p* = 0.624; Figure 2H) were likewise similar between groups.

Total locomotor activity across the recording was lower in cKO mice (Welch’s *t*(14.00) = 7.09, *p* < 0.0001; Figure 2I). Two-way repeated-measures ANOVA of activity averaged within each mouse across the four dark and three light periods identified effects of phase (*F*(1,14) = 232.80, *p* < 0.0001) and genotype (*F*(1,14) = 34.71, *p* < 0.0001), together with a genotype × phase interaction (*F*(1,14) = 58.84, *p* < 0.0001; Figure 2J); activity was lower in cKO mice during the dark phase (Šídák-adjusted *p* < 0.0001) but not the light phase (*p* = 0.504). The hourly activity trace across the full 3.5-day recording is shown in Figure 2K.

Mean respiratory exchange ratio (RER) was lower in cKO mice (Welch’s *t*(8.11) = 5.41, *p* = 0.0006; Figure 2L). RER averaged across the dark and light periods showed effects of phase (*F*(1,14) = 123.09, *p* < 0.0001) and genotype (*F*(1,14) = 34.80, *p* < 0.0001), with no genotype × phase interaction (*F*(1,14) = 0.037, *p* = 0.851; Figure 2M); RER was lower in cKO mice in both the dark and light phases (Šídák-adjusted *p* < 0.0001 for each). The corresponding hourly RER profile is shown in Figure 2N.

Daily mean heat production normalized to lean mass was lower in cKO mice (Welch’s *t*(6.33) = 6.84, *p* = 0.0004; Figure 2O). Dark/light heat production showed effects of phase (*F*(1,14) = 354.00, *p* < 0.0001), genotype (*F*(1,14) = 84.71, *p* < 0.0001), and genotype × phase (*F*(1,14) = 32.34, *p* < 0.0001; Figure 2P), and heat production was lower in cKO mice in both phases (Šídák-adjusted *p* < 0.0001 for each). Hourly heat production over the recording is shown in Figure 2Q.

### Indirect calorimetry during high-fat diet

Daily food intake normalized to lean mass did not differ significantly between genotypes (Welch’s *t*(13.47) = 1.49, *p* = 0.160; Figure 3A). Dark/light food intake showed an effect of phase (*F*(1,14) = 80.77, *p* < 0.0001) and a genotype × phase interaction (*F*(1,14) = 12.86, *p* = 0.00298), whereas the genotype main effect was not significant (*F*(1,14) = 3.04, *p* = 0.103; Figure 3B); cKO mice ate less during the dark phase (Šídák-adjusted *p* = 0.00156) but not the light phase (*p* = 0.366). Cumulative food intake across the recording is shown in Figure 3C.

We next assessed the same measures in the same mice after 3 weeks of HFD (study week 5), again using a 3.5-day metabolic-cage recording. Body weight was lower in cKO mice (Welch’s *t*(13.88) = 2.22, *p* = 0.0432; Figure 3D), whereas lean mass did not differ significantly (Welch’s *t*(13.38) = 1.50, *p* = 0.156; Figure 3E). Percentage lean mass was higher in cKO mice (Welch’s *t*(11.88) = 2.56, *p* = 0.0252; Figure 3F), while fat mass (Welch’s *t*(13.83) = 2.44, *p* = 0.0287; Figure 3G) and percentage fat mass (Welch’s *t*(11.23) = 2.39, *p* = 0.0352; Figure 3H) were lower. Relative body weight after versus before the metabolic-cage recording did not differ significantly between genotypes (Welch’s *t*(13.06) = 1.54, *p* = 0.148; Figure 3I), whereas body-weight gain from the start of HFD was lower in cKO mice (Welch’s *t*(13.01) = 5.52, *p* < 0.0001; Figure 3J).

Total locomotor activity was lower in cKO mice (Welch’s *t*(13.98) = 2.43, *p* = 0.0292; Figure 3K). Two-way repeated-measures ANOVA of average dark- and light-phase activity identified an effect of phase (*F*(1,14) = 50.70, *p* < 0.0001) and a genotype × phase interaction (*F*(1,14) = 7.47, *p* = 0.0162), while the genotype main effect did not reach significance (*F*(1,14) = 4.32, *p* = 0.0567; Figure 3L); activity was lower in cKO mice during the dark phase (Šídák-adjusted *p* = 0.00541) but not the light phase (*p* = 0.998). The hourly activity profile is shown in Figure 3M.

Mean RER did not differ significantly between genotypes (Welch’s *t*(13.83) = 1.68, *p* = 0.116; Figure 3N). Dark/light RER showed a strong phase effect (*F*(1,14) = 207.84, *p* < 0.0001), with no significant genotype effect (*F*(1,14) = 2.37, *p* = 0.146) or genotype × phase interaction (*F*(1,14) = 0.767, *p* = 0.396; Figure 3O). The hourly RER profile is shown in Figure 3P.

Daily mean heat production normalized to lean mass was lower in cKO mice (Welch’s *t*(12.13) = 5.67, *p* < 0.0001; Figure 3Q). Dark/light heat production showed effects of phase (*F*(1,14) = 274.30, *p* < 0.0001) and genotype (*F*(1,14) = 23.56, *p* = 0.0003), together with a genotype × phase interaction (*F*(1,14) = 11.22, *p* = 0.00477; Figure 3R); heat production was lower in cKO mice during both the dark (Šídák-adjusted *p* < 0.0001) and light (*p* = 0.0252) phases. The hourly heat-production trace is shown in Figure 3S.

### Fecal output, energy content, lipids, and microbiome composition

Fecal output was assessed over 5-day periods in male control (*n*=10) and male *Ntsr1^Cre^*;*Th^flox^* cKO (*n*=7) mice used for the feeding analyses (the same mice used in Figure 1A–F). Average daily fecal output on SCD did not differ between genotypes (Welch’s *t*(14.69) = 0.41, *p* = 0.690; Figure 4A). Two-way repeated-measures ANOVA of SCD day/night fecal output identified an effect of phase (*F*(1,15) = 47.30, *p* < 0.0001), with no genotype effect (*F*(1,15) = 1.21, *p* = 0.288) or genotype × phase interaction (*F*(1,15) = 0.127, *p* = 0.727; Figure 4B). On HFD, average daily fecal output was lower in cKO mice (Welch’s *t*(12.96) = 5.38, *p* = 0.0001; Figure 4C). Day/night HFD fecal output showed effects of phase (*F*(1,15) = 6.44, *p* = 0.0227) and genotype (*F*(1,15) = 22.07, *p* = 0.0003), without a significant genotype × phase interaction (*F*(1,15) = 3.73, *p*= 0.0726; Figure 4D); nighttime fecal output was lower in cKO mice (Šídák-adjusted *p* = 0.0005), whereas daytime output did not differ (*p* = 0.607). To test whether altered loss of calories in feces could contribute to obesity resistance on HFD, fecal energy density and fecal mass were assessed under both SCD and HFD conditions (Figure 4E, F). Fecal energy density was higher on HFD than SCD (two-way ANOVA diet effect: *F*(1,42) = 258.8, *p* < 0.0001; Figure 4E) but did not differ between genotypes (*F*(1,42) = 0.278, *p*= 0.601), and there was no genotype × diet interaction (*F*(1,42) = 0.019, *p* = 0.893). Tukey-adjusted comparisons confirmed higher heat of combustion on HFD within both control and cKO mice (both *p* < 0.0001), with no genotype difference on SCD (*p* = 0.965) or HFD (*p* = 0.993). Average fecal mass produced per day likewise did not differ between genotypes on SCD (control 0.90 g, cKO 0.79 g; Welch’s *t*(17.06) = 1.54, *p* = 0.143) or HFD (control 0.20 g, cKO 0.19 g; Welch’s *t*(20.53) = 0.78, *p* = 0.446; Figure 4F). Home-cage food intake, measured in mice from the bomb calorimetry cohort, did not differ between genotypes on SCD (control 11.29 ± 0.79 kcal/day, cKO 10.76 ± 0.50 kcal/day; Welch’s *t*(10.16) = 0.57, *p* = 0.582; Figure 4G) or HFD (control 17.39 ± 0.76 kcal/day, cKO 16.45 ± 1.33 kcal/day; Welch’s *t*(9.53) = 0.61, *p* = 0.553; Figure 4H).

Although bomb calorimetry did not show a genotype difference in fecal energy density, we also measured fecal NEFA concentrations as a complementary measure of fecal lipid composition [32, 33]. Under SCD, fecal NEFA concentration did not differ between genotypes (Welch’s *t*(11.99) = 0.61, *p* = 0.556; Figure 4I). Under HFD, fecal NEFA concentration likewise did not differ significantly between genotypes (Welch’s *t*(9.88) = 0.68, *p* = 0.514; Figure 4J).

Finally, we performed exploratory studies at the genus level of fecal microbiome composition under HFD conditions (Figure 5). Because *Turicibacter* has recently been implicated in host lipid uptake and metabolic health [34], we examined its abundance. The genus-by-genus comparison of mean abundance between groups did not show a difference in microbiome composition between control and cKO mice on HFD (the microbiome was not examined on SCD; *n* = 7 control and *n* = 3 cKO). Overall diversity did not differ between genotypes (Figure 5A-B, Shannon index, mean ± SEM: control 2.95 ± 0.08, cKO 3.02 ± 0.08; Welch’s *t*(6.02) = 0.65, *p* = 0.543). Among the 48 genera with at least 0.1% average abundance, none differed between genotypes after adjusting for multiple comparisons. Here, *q* is the *p* value adjusted for testing 48 genera at once using the Benjamini-Hochberg false-discovery-rate method; the smallest *q* was 0.93, meaning that none remained significant after correction. The largest difference was in *Acetatifactor*, which normally expands in HFD-fed mice [35]. *Acetatifactor* comprised 5.5 ± 1.1% of the microbiome in controls but only 2.3 ± 0.7% in cKO mice (*p* = 0.039, *q* = 0.93). This *p* value is below 0.05 but is not significant after adjustment because, with 48 genera tested, about two would be expected to fall below 0.05 by chance alone. *Dubosiella*, a genus associated with improved lipid metabolism in HFD-fed mice [36], was numerically higher in cKO mice (15.8 ± 6.3%) than in controls (10.2 ± 3.3%; *p* = 0.490, *q* = 0.93). Similarly, *Turicibacter*, the genus implicated in limiting DIO [34], was numerically higher in cKO mice (0.44 ± 0.23%) than in controls (0.02 ± 0.01%), but this difference was not significant (Figure 5C; Welch’s *t*(2.01) = 1.79, *p* = 0.214, *q* = 0.93, ns).

## Discussion

A major unresolved question from our prior work was why loss of dopamine production in *Ntsr1*-expressing neurons protects mice from DIO [23]. Other than a small suppression in day-eating of HFD, we did not detect any other mechanisms by which the *Ntsr1^Cre^*;*Th^flox^*cKO mice could resist DIO. One straightforward possibility was that *Ntsr1^Cre^*;*Th^flox^*cKO mice expend more energy through increased thermogenesis. The present data do not support that model. Across independent measures, cKO mice showed lower heat production, reduced locomotor activity, and lower subcutaneous temperature. Thus, DIO resistance occurred despite a physiological profile that would generally be expected to conserve energy and favor, rather than prevent, weight gain.

Instead, reduced HFD intake is a likely proximal contributor to the phenotype in the present cohort. In our prior study, cKO mice were slightly hyperphagic on SCD and consumed similar amounts of HFD as controls [23]. Here, cKO mice consumed less HFD but not SCD (Figure 1C-F) in some experiments whereas in others the food intake was similar on both SCD and HFD (Figure 4G-H). A modest daily difference in HFD intake could accumulate over several weeks and contribute substantially to the reduced weight gain. Upon transitioning to the metabolic cages, cKO mice decreased feeding on SCD dramatically (Figure 2A), but maintained normal feeing on HFD (Figure 3A). Thus, the present data support reduced HFD intake in some contexts as a contributor to the DIO-resistance of *Ntsr1^Cre^*;*Th^flox^* cKO male mice, but overall we cannot say that the cKO mice are clearly hypophagic.

We did not detect genotype differences in the fecal endpoints used to assess energy or lipid loss. Fecal output was assessed in two independent cohorts. Feces from the first cohort were collected exclusively for bomb calorimetry (Figure 4E, F), whereas feces from the second cohort were collected exclusively for measurement of NEFA (Figure 4A-D). In the bomb calorimetry cohort, fecal output did not differ between genotypes on either SCD or HFD. Food intake in this cohort likewise did not differ between genotypes on SCD or HFD (Figure 4G, H). In the NEFA cohort, cKO mice produced less fecal material than controls on HFD, but not on SCD (Figure 4A,C). Thus, the reduction in fecal output observed in HFD-fed cKO mice was cohort-specific and was not consistently observed across the two experiments. The lower HFD fecal output of the NEFA cohort, which was drawn from the replication cohort, parallels the reduced HFD intake of those mice (Figure 1E), whereas the bomb calorimetry cohort showed neither reduced HFD intake nor reduced fecal output. This pattern suggests that fecal output tracked food intake rather than absorption, and that the HFD hypophagia of cKO mice was not evident in every cohort.

Fecal energy density and fecal NEFA concentrations did not differ between genotypes. Bomb calorimetry measures total combustible energy, whereas direct assessment of dietary fat absorption requires lipid-specific methods or a complete fat-balance [32, 33]. An additional untested possibility is that cKO mice lose lipid through the skin as sebum, a mechanism shown to cause adipose loss in mice without changes in food intake, fecal energy loss, or energy expenditure [37]. Because we did not assess coat lipid content, sebaceous gland morphology, or sebum-associated gene expression, this possibility remains speculative.

Taken together, these findings indicate that resistance to HFD-induced weight gain in *Ntsr1^Cre^*;*Th^flox^* cKO mice is not fully explained by measured components of energy balance. Reduced HFD intake is the most likely proximal contributor in the present cohort, although its causal relationship to lower body weight remains unresolved and the phenotype does not consistently reproduce the previously observed daytime-feeding effect [23]. The exploratory microbiome analysis did not identify a genotype-level difference, although *Turicibacter* and *Dubosiella* were numerically more abundant in cKO mice. The causal relevance of these taxa in the present model therefore remains unknown.

## CRediT authorship contribution statement

RW: Writing – review & editing, Writing – original draft, Visualization, Validation, Project administration, Methodology, Investigation, Formal analysis, Data curation, and Conceptualization.

AG – Methodology, Investigation, Formal analysis, Data curation

JTW – Methodology, Investigation, Data curation.

RT – Methodology, Investigation, Formal analysis, Data curation.

AKK – Investigation, Formal analysis, Data curation.

ADS – Writing – review & editing, Writing – original draft, Supervision, Resources, Project administration, Formal analysis, Data curation, and Conceptualization.

## Acknowledgment

Funding was provided by the National Institute of General Medical Sciences of the National Institutes of Health under award number R16GM145576, National Institute of Diabetes and Digestive and Kidney Diseases under award number DK-132736. The content is solely the responsibility of the authors and does not necessarily represent the official views of the National Institutes of Health. The Metabolic Phenotyping Core is partially supported by funding from the State of Texas and NIH NORC grant 1P30DK127984-01A1. The authors Thanks UT Southwestern Metabolic Phenotyping Core (RRID:SCR_026404) for the analysis of fecal energy content (bomb calorimetry) and fatty acids.

## Supplementary materials

There is one supplemental Figure associated with this manuscript.

## Data availability

Data will be made available on request.

## Declaration of generative AI and AI-assisted technologies in the manuscript preparation process

During the preparation of this work the author(s) used ChatGPT and Claude to edit writing. After using this tool/service, the authors reviewed and edited the content as needed and take full responsibility for the content of the published article.

---

**Supplementary Figure 1.** Mouse cohorts used in this study. Each mouse is listed with its ID, sex, *Ntsr1^Cre^*and *Th* genotype, and experimental group. Cohort 1 (replication; control *n* = 10, cKO *n* = 7) was used for body weight, home-cage food intake, and fecal output (Figure 1A–F, Figure 4A–D), with subsets used for fecal NEFA (Figure 4I, J) and microbiome analysis (Figure 5). Cohort 2 (control *n* = 10, cKO *n* = 6) was used for indirect calorimetry and body composition (Figures 2 and 3). Cohort 3 (control *n* = 13, cKO *n* = 10) was used for bomb calorimetry (Figure 4E, F), and 7 control and 7 cKO mice from this cohort were also used for home-cage food intake (Figure 4G, H). Cohort 4 (control *n* = 6, cKO *n* = 4) was used for Nanotag temperature and activity (Figure 1G–N). All mice were male.

