## Supplemental Figure 1 for "Tyrosine hydroxylase deletion in *Ntsr1* neurons confers resistance to diet-induced weight gain: an energy-balance analysis"

### Supplementary Figure 1

##### Cohort 1: Replication (control n = 10, cKO n = 7)

*Body weight, home-cage food intake and fecal output (Figure 1A–F, 4A–D); fecal NEFA (Figure 4I, J) and microbiome (Figure 5) in subsets*

| Mouse ID | Sex | *Ntsr1*$^{Cre}$ | *Th* | Group |
|---|---|---|---|---|
| 925 | M | + | flox/wt | Control |
| 926 | M | − | wt/wt | Control |
| 930 | M | + | flox/wt | Control |
| 943 | M | + | wt/wt | Control |
| 945 | M | + | flox/wt | Control |
| 961 | M | − | wt/wt | Control |
| 965 | M | + | wt/wt | Control |
| 966 | M | − | flox/wt | Control |
| 967 | M | + | wt/wt | Control |
| 968 | M | − | flox/wt | Control |
| 927 | M | + | flox/flox | cKO |
| 928 | M | + | flox/flox | cKO |
| 941 | M | + | flox/flox | cKO |
| 962 | M | + | flox/flox | cKO |
| 963 | M | + | flox/flox | cKO |
| 964 | M | + | flox/flox | cKO |
| 969 | M | + | flox/flox | cKO |

##### Cohort 2: Metabolic cages (control n = 10, cKO n = 6)

*Indirect calorimetry and body composition (Figures 2 and 3)*

| Mouse ID | Sex | *Ntsr1*$^{Cre}$ | *Th* | Group |
|---|---|---|---|---|
| 753 | M | − | flox/wt | Control |
| 754 | M | − | flox/wt | Control |
| 755 | M | + | flox/wt | Control |
| 756 | M | − | flox/wt | Control |
| 757 | M | + | wt/wt | Control |
| 763 | M | − | flox/flox | Control |
| 770 | M | − | flox/wt | Control |
| 771 | M | − | flox/flox | Control |
| 779 | M | + | wt/wt | Control |
| 780 | M | + | flox/wt | Control |
| 744 | M | + | flox/flox | cKO |
| 752 | M | + | flox/flox | cKO |
| 764 | M | + | flox/flox | cKO |
| 765 | M | + | flox/flox | cKO |
| 766 | M | + | flox/flox | cKO |
| 769 | M | + | flox/flox | cKO |

##### Cohort 3: Bomb calorimetry (control n = 13, cKO n = 10)

*Fecal energy and mass (Figure 4E, F); home-cage food intake in a subset (Figure 4G, H; control n = 7, cKO n = 7)*

| Mouse ID | Sex | *Ntsr1*$^{Cre}$ | *Th* | Group |
|---|---|---|---|---|
| *Bomb calorimetry only (feces collected February 2025)* | | | | |
| 795 | M | − | flox/wt | Control |
| 798 | M | − | flox/wt | Control |
| 808 | M | − | flox/wt | Control |
| 819 | M | + | flox/wt | Control |
| 820 | M | − | flox/flox | Control |
| 821 | M | − | flox/flox | Control |
| 809 | M | − | flox/wt | cKO |
| 817 | M | + | flox/flox | cKO |
| 818 | M | + | flox/flox | cKO |
| *Bomb calorimetry and home-cage food intake* | | | | |
| 829 | M | − | flox/wt | Control |
| 846 | M | − | flox/wt | Control |
| 847 | M | + | flox/wt | Control |
| 851 | M | + | flox/wt | Control |
| 852 | M | + | flox/wt | Control |
| 854 | M | − | flox/flox | Control |
| 861 | M | − | flox/flox | Control |
| 828 | M | + | flox/flox | cKO |
| 848 | M | + | flox/flox | cKO |
| 849 | M | + | flox/flox | cKO |
| 850 | M | + | flox/flox | cKO |
| 853 | M | + | flox/flox | cKO |
| 859 | M | + | flox/flox | cKO |
| 860 | M | + | flox/flox | cKO |

##### Cohort 4: Nanotag (control n = 6, cKO n = 4)

*Subcutaneous temperature and activity (Figure 1G–N)*

| Mouse ID | Sex | *Ntsr1*$^{Cre}$ | *Th* | Group |
|---|---|---|---|---|
| 681 | M | + | flox/wt | Control |
| 693 | M | − | flox/flox | Control |
| 694 | M | − | flox/flox | Control |
| 702 | M | − | flox/flox | Control |
| 707 | M | − | flox/flox | Control |
| 708 | M | − | flox/flox | Control |
| 691 | M | + | flox/flox | cKO |
| 692 | M | + | flox/flox | cKO |
| 701 | M | + | flox/flox | cKO |
| 706 | M | + | flox/flox | cKO |
